# Correcting reference bias from the Illumina Isaac aligner enables analysis of cancer genomes

**DOI:** 10.1101/836171

**Authors:** Alex J. Cornish, Daniel Chubb, Anna Frangou, Phuc H. Hoang, Martin Kaiser, David C. Wedge, Richard S. Houlston

## Abstract

Estimating the fraction of cancer cells with individual somatic mutations is central to many analyses in cancer genomics, including characterisation of clonal architecture and timing of mutational events. Estimation of these cancer cell fractions (CCFs) is contingent on unbiased assessment of the fraction of reads supporting variant alleles (VAFs). We demonstrate that VAFs computed by the Illumina Isaac pipeline, used in many large-scale sequencing projects including The 100,000 Genomes Project, are biased by the preferential soft clipping of reads supporting non-reference alleles (semi-aligned reads). We show that these biased VAFs can have deleterious effects on downstream analyses reliant on unbiased CCF estimates. While Isaac bias can be corrected through realignment with alternative parameters, this is computationally intensive. We therefore developed FixVAF, a tool for removing bias introduced by soft clipping of semi-aligned reads, facilitating downstream analyses without the need for realignment. FixVAF is freely available at https://github.com/danchubb/FixVAF.

## 1. INTRODUCTION

The Illumina Isaac pipeline (Raczy et al., 2013) has been used in a number of large-scale sequencing projects, most notably The 100,000 Genomes Project (100KGP), a £300 million endeavor tasked with transforming genomic research and clinical application in the UK (Klintman et al., 2018; Turnbull et al., 2018). Additionally, the Isaac pipeline has been employed in numerous cancer-sequencing studies (Burns et al., 2018; Quigley et al., 2018; Ross-Innes et al., 2015), to evaluate genome editing and characterise off-target effects (Kim et al., 2015; Kim et al., 2017; Ma et al., 2017), and in the validation of embryo screening techniques (Fiorentino et al., 2014). The pipeline was developed primarily as a faster alternative to approaches such as the community-standard combination of Burrows-Wheeler Alignment (BWA) (Li and Durbin, 2009) and the Genome Analysis Tool Kit (GATK) (McKenna et al., 2010). Whilst the Isaac pipeline has been benchmarked with respect to variant calling (Klintman et al., 2018; Mainzer et al., 2015; Raczy et al., 2013), there has been less extensive evaluation of its suitability for other analyses routine in genomic research, particularly those applied in cancer genomics.

Many analyses that use tumour sequencing data, including the characterisation of clonal architecture (Dentro et al., 2018; Jamal-Hanjani et al., 2017; Nik-Zainal et al., 2012; Williams et al., 2018), assessment of cancer drivers and mutational processes (McGranahan et al., 2015), and timing of mutational events (Gerstung et al., 2018; Mitchell et al., 2018) are reliant on an unbiased estimation of the proportion of cancer cells containing individual mutations. Such cancer cell fractions (CCFs) are computed using the fraction of aligned reads supporting the variant allele (VAF), the tumour copy number profile and the tumour sample purity (Dentro et al., 2017). Any bias in the alignment of reads, favoring either the reference or non-reference (alternate) allele, could therefore bias CCF estimates and negatively impact analyses.

We assessed the Isaac pipeline to determine its suitability for analyses reliant on CCF estimation. After demonstrating that the soft clipping of semi-aligned reads performed by the Isaac aligner introduces reference bias, we characterise potential negative effects of this bias on downstream analyses, including calling of copy number variants (CNVs), purity estimation and clonality reconstruction. To address this shortcoming we developed FixVAF, a tool that removes sources of bias from VAFs computed from Isaac sequence alignments, facilitating robust CCF estimation.

## 2. MATERIALS AND METHODS

### 2.1. Sequencing data

Whole genome sequencing (WGS) data were obtained from 25 tumour-normal multiple myeloma (MM) pairs from the UK National Cancer Research Institute Myeloma XI trial (Jackson et al., 2019). Subjects were included on the basis of DNA sample availability, and informed consent was obtained from all patients. Tumour DNA was extracted from plasma cells selected and sorted using CD138 micro beads (Walker et al., 2010), whilst germline DNA was derived from matched blood samples. Sequencing libraries were prepared using Illumina SeqLab specific TruSeq Nano High Throughput library preparation kit (Illumina Inc., San Diego, CA 92122 USA) and Illumina HiSeqX technology was used to conduct paired end sequencing. FastQC (v.0.11.4) was used to quality check raw WGS data. The Myeloma XI trial was approved by the Oxfordshire Research Ethics Committee (MREC 17/09/09, ISRCTN49407852) and conducted in accordance with the Declaration of Helsinki and Good Clinical Practice.

### 2.2. Sequence alignment and variant calling

The Isaac aligner (Raczy et al., 2013) has a parameter (--clip-semialigned) that invokes the soft clipping of reads at each end until a stretch of five consecutive bases are matched with the reference sequence. When referring to ‘soft clipping of semi-aligned reads’, we are exclusively referring to soft clipping invoked by the --clip-semialigned parameter, as Isaac also soft clips reads for other reasons. This parameter is present in all Isaac versions after v01.13.06.20 (**Supplementary Table 1**) and was used in the alignment of 100KGP data.

Sequencing data were aligned to the *Homo sapiens* GRCh38Decoy assembly using two approaches: (1) Isaac v03.16.02.19 with soft clipping of semi-aligned reads (--clip-semialigned=1) and (2) Isaac v03.16.02.19 without soft clipping of semi-aligned reads (--clip-semialigned=0) (**Figure 1**). Illumina provided sequence alignments generated with soft clipping of semi-aligned reads enabled for the 25 tumour-normal pairs as part of their sequencing services. We subsequently realigned these data using Isaac without soft clipping of semi-aligned reads. Isaac v03.16.02.19 was considered as this software version was used in 100KGP and therefore of particular interest.

**Figure 1:**
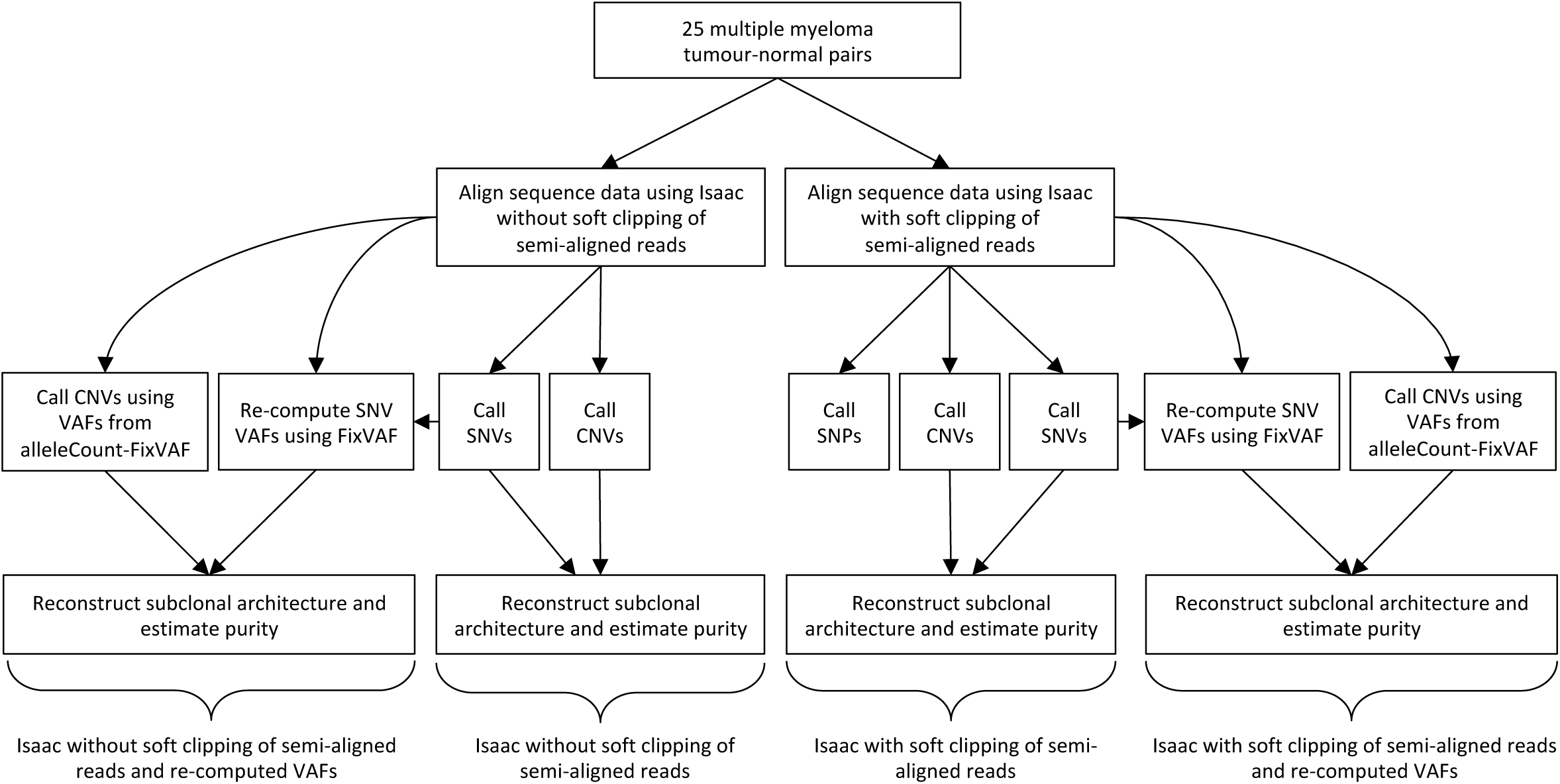
Data processing flowchart. Single nucleotide polymorphisms (SNPs), single nucleotide variants (SNVs) and copy number variants (CNVs) called using Starling, Strelka and Battenberg respectively. Subclonal architectures were reconstructed using DPClust and tumour sample purity was re-estimated using Ccube. VAF: variant allele frequency.

Germline variants were called by Illumina using Starling v2.4.7 (Raczy et al., 2013). Germline single nucleotide polymorphism (SNP) VAFs were calculated directly from Binary Sequence Alignment Map (BAM) files using alleleCount with minimum base and mapping qualities of 20 and 35 respectively (Van Loo et al., 2010). Somatic single nucleotide variants (SNVs) were called using Strelka v2.4.7 (Kim et al., 2018; Saunders et al., 2012). Parameters used to run Isaac, Starling and Strelka are detailed in **Supplementary Table 2**. Differences in VAF distributions were assessed using Wilcoxon Signed-Rank tests.

### 2.3. Removing bias introduced by soft clipping of semi-aligned reads

The Isaac --clip-semialigned parameter invokes the soft clipping of read ends until five consecutive bases are matched with the reference genome (Illumina, 2016). This soft clipping therefore results in the loss of support for alternate alleles occurring within five bases of each read end. To reduce allelic bias introduced by this clipping, FixVAF soft clips all reads by five bases at each end, regardless of whether any of the bases are variant sites or whether the reads support reference or alternate alleles (**Figures 2** and **3a**). Reads containing small insertions and deletions at variant positions are ignored. FixVAF can be applied to Strelka and Starling output, requires the pysam library, and is available as a Python3 package from GitHub (https://github.com/danchubb/FixVAF). Required inputs are a BAM file and a Variant Call Format (VCF) file. Counts of all four possible alleles at each position are provided in an additional INFO field. Strelka provides two tiers of counts, T1 and T2, requiring read mapping qualities of ≥40 and ≥5 respectively. FixVAF computes unbiased counts using these two quality thresholds, along with a third tier, T3, requiring a read mapping quality >0. Providing counts using different quality thresholds allows users to choose how conservative they wish the read filtering to be.

**Figure 2:**
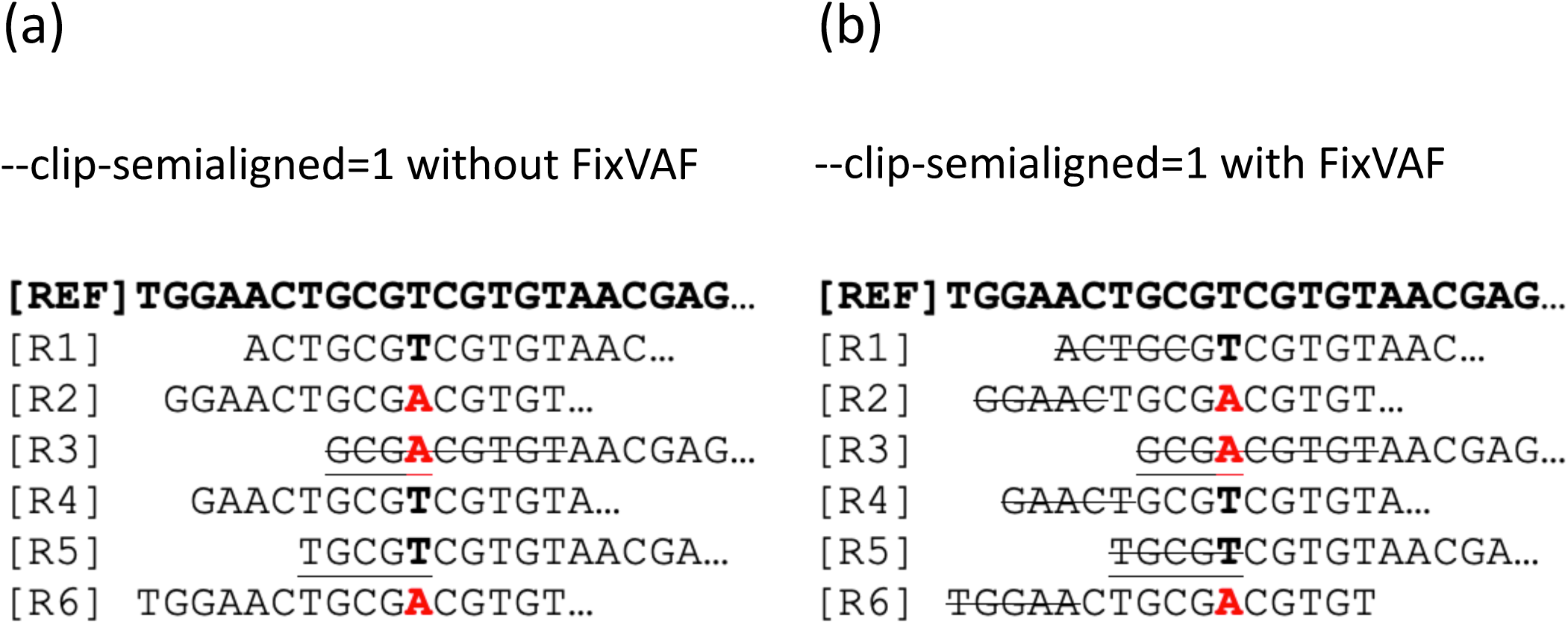
Removing bias introduced by soft clipping of semi-aligned reads. Alignment of six reads to a reference sequence containing an A/T variant. Bold black T and red A represent reference and alternate alleles respectively. Soft clipping is represented by strikethrough. Without soft clipping, three reads would support both the reference (T) and alternate (A) alleles, resulting in an unbiased variant allele frequency (VAF) of 3/6=0.5. (**a**) Read R3 is soft clipped until five consecutive matches with the reference are obtained. After clipping, only two reads support the alternate allele (A), whilst three reads support the reference allele (T), resulting in a biased VAF of 2/5=0.4. (**b**) FixVAF clips all reads by five bases, regardless of whether they contain a variant site or support a reference or alternate allele. Reads supporting both the reference and alternate alleles are now clipped by five bases. In this example, FixVAF would compute a VAF of 2/4=0.5, and therefore remove bias.

**Figure 3:**
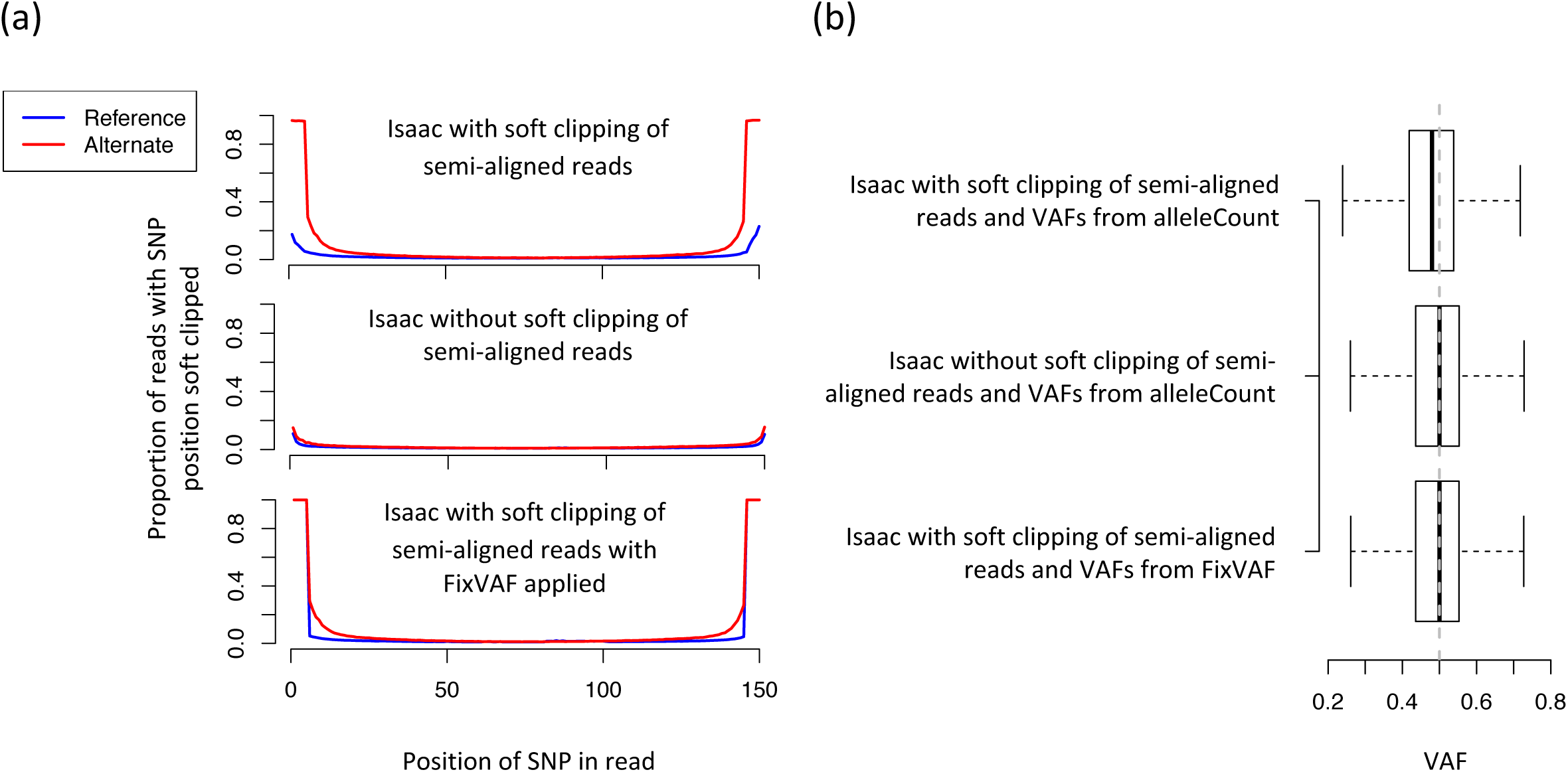
Evidence of reference bias from sequence aligners. (**a**) Proportion of reads covering single-nucleotide polymorphism (SNP) positions supporting the reference (blue line) and alternate (red line) alleles with that read position soft clipped. (**b**) SNP variant allele frequency (VAF) distributions in whole genome data from 25 normal samples. Sequencing data were aligned using Isaac with and without soft clipping of semi-aligned reads. VAFs were computed from Binary Sequence Alignment Map files using alleleCount or FixVAF. Grey dashed line represents expected median VAF of 0.5. VAFs computed as the number of reads supporting the alternate allele divided by the number of reads supporting either the alternate or reference alleles. Whiskers extend 1.5 times inter-quartile range and values outside of this range are not shown.

### 2.4. Calling and comparing copy number profiles

Reconstruction of clonal and subclonal CNVs was conducted using Battenberg v2.2.8 (Nik-Zainal et al., 2012). Battenberg computes VAFs using alleleCount, and to remove bias from these VAFs we therefore modified alleleCount to ignore positions within five bases of each read end, as per FixVAF. This modified version of alleleCount, which we refer to as alleleCount-FixVAF, is available from GitHub (https://github.com/danchubb/alleleCount-FixVAF).

Battenberg phases variants using IMPUTE2 (Howie et al., 2012), which is implemented for hg37. To call CNVs, SNP positions were therefore converted to hg37 before running Battenberg and output segment positions were converted back to hg38. CNVs were called using Isaac alignments generated both with and without soft clipping of semi-aligned reads, and VAFs computed using alleleCount and alleleCount-FixVAF (**Figure 1**). Battenberg was run with the IMPUTE2 seed variable set to 0.

We compared copy number profiles from the same samples to determine whether soft clipping of semi-aligned reads affects Battenberg. We define the state of a genomic region (one or more contiguous bases) to be different between two copy number profiles if (i) subclonal changes are not present in either profile and clonal copy number states differ, (ii) subclonal changes are present in both profiles and subclonal copy number states differ, or (iii) a subclonal change is present in one profile but not the other (**Figure 4b**). Comparisons are not limited to discreet segments of continuous copy number called by Battenberg, but are instead conducted base-by-base.

**Figure 4:**
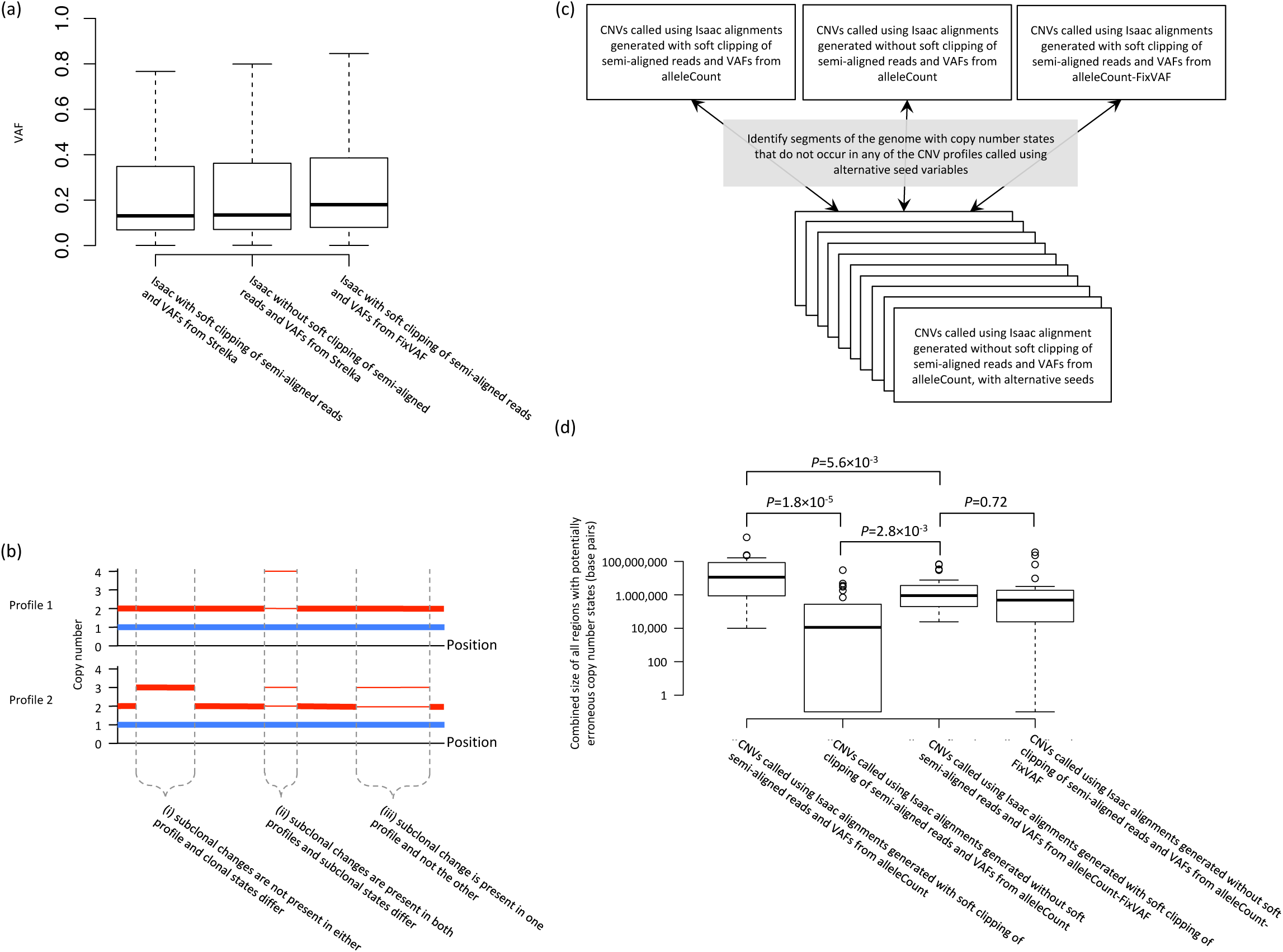
Effects of soft clipping of semi-aligned reads on somatic variants. (**a**) Single nucleotide variant (SNV) variant allele frequencies (VAFs) from 25 tumour-normal pairs aligned with and without soft clipping of semi-aligned reads. Only SNVs identified using both alignments were considered. Whiskers extend 1.5 times inter-quartile range and values outside this range are not shown. Distribution differences assessed using Wilcoxon Signed-Rank tests, and for all comparisons *P*<2.2×10^−16^. (**b**) Identifying differences between copy number profiles. The copy number state of a genomic region is defined as different if (i) subclonal changes are not present in either profile and clonal states differ, (ii) subclonal changes are present in both profiles and subclonal states differ, or (iii) a subclonal change is present in one profile and not the other. Red and blue lines represent total and minor copy number states respectively. Thick and thin horizontal lines represent clonal and subclonal states respectively. Dashed lines demark regions with states that differ between the two profiles. (**c**) Strategy for assessing whether soft clipping of semi-aligned reads affects Battenberg, and how application of alleleCount-FixVAF influences such effect. (**d**) Total size of potentially erroneous copy number state calls from profiles generated from each alignment, with VAFs computed using alleleCount and alleleCount-FixVAF. Distribution differences assessed using Wilcoxon Signed-Rank tests. CNV: copy number variant.

Parts of the Battenberg algorithm, including variant phasing by IMPUTE2 (Howie et al., 2012; Nik-Zainal et al., 2012), are stochastic, meaning that different copy number profiles are output depending on the seed variable specified. To evaluate whether differences between copy number profiles from alignments generated with and without soft clipping of semi-aligned reads are greater than differences between profiles generated using different IMPUTE2 seed variables, we compared profiles generated using IMPUTE2 seeds of 0 to profiles generated from alignments without soft clipping of semi-aligned reads with ten alternative IMPUTE2 seed variables (ranging from 1 to 10). If the state of a genome region called using an IMPUTE2 seed of 0 is never observed in any of the profiles generated using alternative seed variables, then we consider the state of the region potentially erroneous. Such discrepancies could occur if multiple copy number states explain the observed allele frequencies similarly well, or be due to the soft clipping of semi-aligned reads. By comparing the total size of potentially erroneous regions between alignments generated with and without soft clipping of semi-aligned reads, and with and without correction of VAF bias, we assess firstly whether soft clipping of semi-aligned reads affects Battenberg, and secondly whether computation of unbiased VAFs reduces such effect.

### 2.5. Assessing clonality

Subclonal reconstruction was conducted with DPClust v2.2.8 (Nik-Zainal et al., 2012) using SNVs from autosomes and the X chromosome. Clusters of mutations containing <1% of all mutations were excluded. DPClust was run using SNVs and copy number profiles generated using sequence alignments with and without soft clipping of semi-aligned reads and VAFs from Strelka, and again using sequencing alignments with soft-clipping of semi-aligned reads but with VAFs from FixVAF (**Figure 1**). Putative clonal mutation clusters were defined in each sample as the cluster with a CCF closest to 1.

### 2.6. Tumour sample purity re-estimation

We re-estimated tumour sample purity using Ccube v1.0 (Yuan et al., 2018) to assess the effect of soft clipping of semi-aligned reads on purity estimation. Ccube requires as input whether or not a tumour has undergone whole genome duplication, and we therefore considered whole genome duplication to have occurred if the Battenberg ploidy estimate was >2.6. Ccube was run using copy number profiles generated using sequence alignments with and without soft clipping of semi-aligned reads and autosomal SNV VAFs from Strelka, and again using sequencing alignments with soft clipping of semi-aligned reads but with autosomal SNV VAFs from FixVAF (**Figure 1**).

### 2.7. Software and data availability

Isaac (v03.16.02.19), Strelka (v2.4.7), alleleCount (v4.0.0), Battenberg (v2.2.8), DPClust (v2.2.8) and Ccube (v1.0) were downloaded from GitHub (**Supplementary Table 2**). *Homo sapiens* GRCh38Decoy reference assembly was downloaded from Illumina (**Supplementary Table 2**). FixVAF and alleleCount-FixVAF are available to download from GitHub (https://github.com/danchubb/FixVAF and https://github.com/danchubb/alleleCount-FixVAF respectively). Analyses were conducted using R versions 3.3.1 and 3.5.0.

## 3. RESULTS

### 3.1. Soft clipping of semi-aligned reads introduces reference bias

In normal tissue samples, sequencing of reads supporting heterozygous SNP reference and alternate alleles can be expected to occur with equal probability. However, due to limited sequencing depth, the exact number of reads supporting reference and alternate alleles will not be equal in many instances, even when the aligner is unbiased. It is nevertheless reasonable to expect that the median VAF of a large number of heterozygous SNPs will approximate 0.5. To evaluate the Isaac aligner, we therefore assessed heterozygous SNP VAF distributions in sequencing data from 25 normal tissue samples (**Figure 3b**; **Supplementary Table 3**), aligned by Isaac using the same parameters employed by 100KGP. Median VAFs per sample ranged from 0.478 to 0.479. This directionally consistent deviation from 0.5 indicates that Isaac can exhibit bias towards the reference allele.

Soft clipping of semi-aligned reads was invoked to generate sequence alignments for the 100KGP. To test whether this clipping contributes reference bias, we realigned the sequencing data from the normal samples using Isaac without soft clipping semi-aligned reads. When semi-aligned reads were not soft clipped, the median VAF of heterozygous SNPs in each sample equaled 0.500 (**Figure 3b**), demonstrating that the clipping introduces reference bias. Soft clipping of semi-aligned reads results in the clipping of the majority of reads supporting the alternate allele where the variant position is within five bases of either read end (**Figure 3a**). Fewer reads supporting the reference allele are soft clipped (**Figure 3a**) and VAFs therefore become biased towards the reference allele. Such disparity in clipping rates between reads supporting reference and alternate alleles is not observed when semi-aligned reads are not soft clipped (**Figure 3a**).

Unlike SNPs in germline samples, we do not know the true VAF of somatic SNVs in tumour samples due to copy number variation, normal sample contamination and clonal heterogeneity. However, SNV VAFs from alignments generated with soft clipping of semi-aligned reads were lower than SNV VAFs from the same samples from alignments generated without soft clipping of semi-aligned reads (*P*<2.2×10^−16^; **Figure 4a**), indicating that this clipping also affects SNV VAFs.

### 3.2. FixVAF removes bias introduced by soft clipping of semi-aligned reads

Whilst SNP VAFs from germline sample sequence alignments generated with soft clipping of semi-aligned reads exhibited reference bias (median VAFs ranged from 0.478 to 0.479 per sample), VAFs computed using FixVAF from the same alignments did not exhibit bias (median VAFs equaled 0.500 for all samples; **Figure 3b**). Furthermore, in tumour samples, SNV VAFs computed using FixVAF from alignments generated with soft clipping of semi-aligned reads were greater than SNV VAFs from Strelka for the same alignments (*P*<2.2×10^−16^; **Figure 4a**). FixVAF SNV VAFs from alignments generated with soft clipping of semi-aligned reads were also greater than Strelka SNV VAFs from alignments generated without soft clipping of semi-aligned reads (*P*<2.2×10^−16^; **Figure 4a**), suggesting the Strelka algorithm may be a source of additional bias.

### 3.3. Soft clipping of semi-aligned reads affects downstream analyses

To assess whether soft clipping of semi-aligned reads affects Battenberg, we compared copy number profiles generated using alignments with and without soft clipping of semi-aligned reads to profiles generated with alternative IMPUTE2 seed values, to identify regions of the genome with potentially erroneous copy number state calls (**Figures 4b** and **4c**). The total size of genome regions with potentially erroneous state calls was greater when Battenberg was run using alignments generated with soft clipping of semi-aligned reads than without (*P*=1.8×10^−5^; **Figure 4d**), demonstrating that soft clipping of semi-aligned reads affects Battenberg. Applying alleleCount-FixVAF to alignments generated with soft clipping of semi-aligned reads reduced the total size of regions with potentially erroneous states >10-fold (*P*=5.6×10^−3^; **Figure 4d**), thereby addressing the bias introduced by clipping. Whilst the total size of these regions was greater than in profiles generated without soft clipping of semi-aligned reads (*P*=2.8×10^−3^; **Figure 4d**), no significant difference was observed between the total size of potentially erroneous regions when alleleCount-FixVAF was applied to alignments generated with and without soft clipping of semi-aligned reads (*P*=0.72; **Figure 4b**). Therefore, whilst the reduction in effective sequencing depth invoked by alleleCount-FixVAF may affect some copy number calls, these observations are consistent with alleleCount-FixVAF removing bias introduced by clipping.

We ran DPClust using alignments generated with and without soft clipping of semi-aligned reads to test the effect of the clipping on subclonal reconstruction. If SNV VAFs do not exhibit allelic bias, we would expect DPClust to identify clusters of mutations with CCFs centered on 1, representing clonal mutations. When DPClust was run using alignments generated with soft clipping of semi-aligned reads, the median CCF of putative clonal mutation clusters in the 25 tumours was 0.959 (**Figure 5a**). Conversely, when DPClust was run using alignments generated without soft clipping of semi-aligned reads, the median CCF of putative clonal mutation clusters was 0.983 (**Figure 5a**), demonstrating that read soft clipping affects clonal mutation characterisation. When DPClust was run using FixVAF VAFs from alignments generated with soft clipping of semi-aligned reads, the median CCF of putative clonal mutation clusters was 0.989 (**Figure 5a**), consistent with FixVAF substantially reducing the effect of bias on clonality reconstruction.

**Figure 5:**
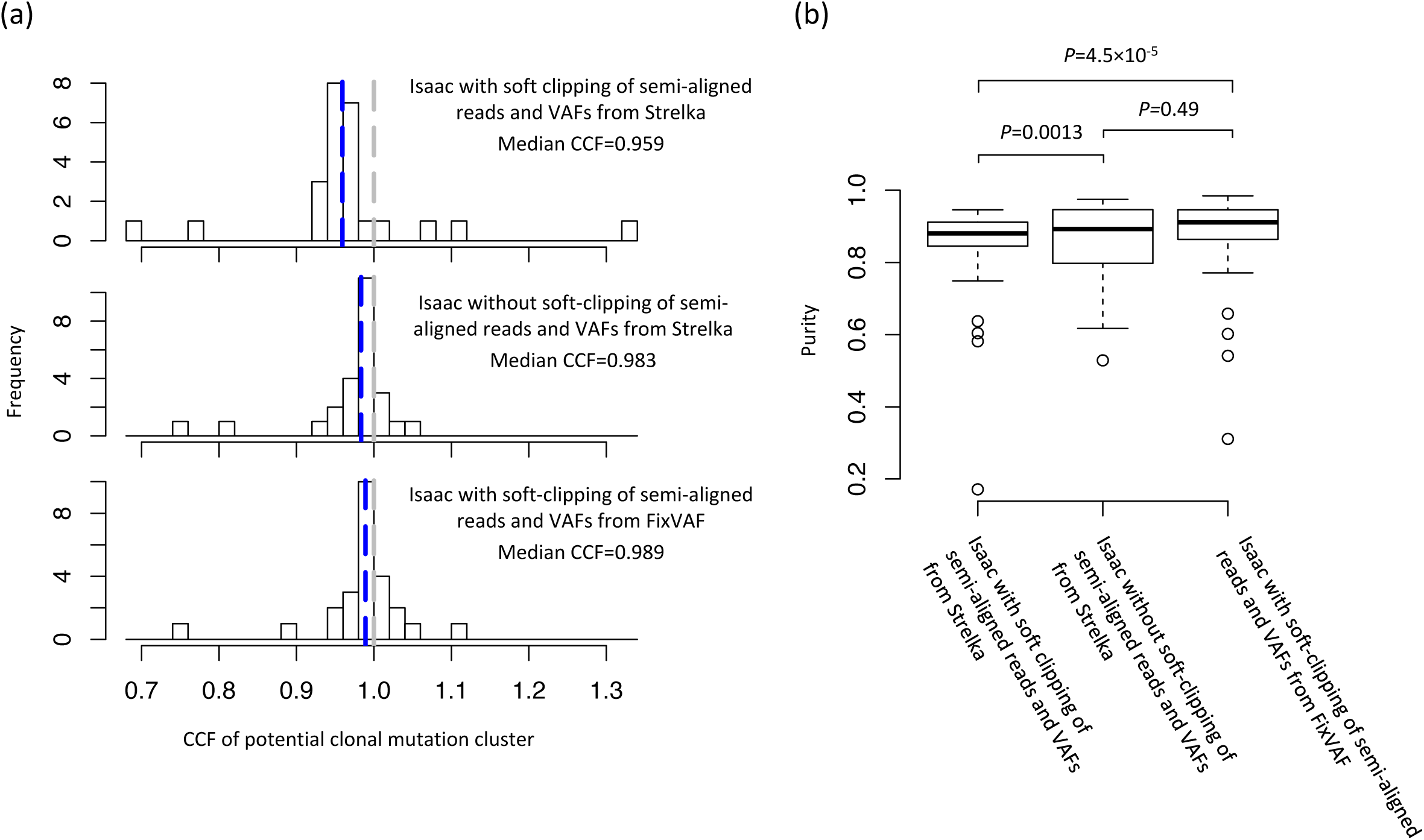
Effect of soft clipping of semi-aligned reads on downstream analyses. (**a**) Cancer cell fractions (CCFs) of clonal mutation clusters identified by DPClust when run with Strelka variant allele frequencies (VAFs) from alignments generated with and without soft clipping of semi-aligned reads, and FixVAF VAFs from alignments generated with soft clipping of semi-aligned reads. Putative clonal mutation clusters were defined in each sample as the cluster with a CCF closest to 1. Blue and grey dashed lines denote the median putative clonal mutation cluster CCF and a CCF of 1 respectively. (**b**) Ccube sample purity estimates computed using Strelka VAFs from alignments generated with and without soft clipping of semi-aligned reads, and using FixVAF VAFs from alignments generated with soft clipping of semi-aligned reads. Whiskers extend up to 1.5 times inter-quartile range. Distribution differences assessed using Wilcoxon Signed-Rank tests.

Ccube tumour sample purity estimates were smaller when computed using alignments generated with soft clipping of semi-aligned reads than without (*P*=1.3×10^−3^; **Figure 5b**), demonstrating that this clipping also affects purity estimation. Purities estimated using FixVAF VAFs from alignments generated with soft clipping of semi-aligned reads were not significantly different from those estimated from non-soft-clipped data (*P*=0.49; **Figure 5b**), consistent with FixVAF also reducing the effect of bias on purity estimation.

## 4. DISCUSSION

Genomic analyses of tumors are becoming increasingly sophisticated, allowing investigation of their histories and the mutations and processes that drive their development (Jolly and Van Loo, 2018; Turajlic et al., 2019). However, these analyses place many assumptions on the data they use (Dentro et al., 2017). To ensure these assumptions are valid, extensive benchmarking of pipelines that process data is required. Otherwise, as the IBM programmer Wilf Hey is accredited with saying: “Garbage in, garbage out” (Elliott et al., 2006).

Soft clipping of semi-aligned reads performed by the Isaac aligner introduces bias towards the reference allele. Such bias can affect downstream processes, potentially making conclusions unreliable for many types of cancer analysis. While the Isaac aligner version assessed in this study (v03.16.02.19) was released in April 2016, as of November 2019 it is still being used to generate sequence alignments in projects such as 100KGP. Whether reference bias exhibited by the Isaac pipeline has affected studies using these data is difficult to predict. It is essential that aligners, such as Isaac, be evaluated to ensure that the data they produce are not systematically biased.

If unbiased VAFs are required, Isaac should be run with soft clipping of semi-aligned reads disabled, or an alternative aligner such as BWA (Li and Durbin, 2009) should be used. Although realignment can be performed where clipped alignments have been previously produced, this may be cost or time-prohibited. For example, projects such as 100KGP have already sequenced and aligned >10,000 tumour-normal genome pairs (Genomics England, 2019). FixVAF computes unbiased VAFs from biased Isaac alignments, thereby enabling downstream analyses reliant on unbiased VAFs without the need for sequencing data realignment.

FixVAF has a number of limitations. Firstly, it is only able to compute VAFs for SNPs and SNVs. Estimating VAFs of small insertions and deletions is more complex, as it can require realignment of reads at the site of the variant (Saunders et al., 2012). Secondly, some variant callers, including Strelka, perform a number of additional read filtering steps (Kim et al., 2018; Saunders et al., 2012) and VAFs computed by FixVAF will therefore not necessarily equal VAFs computed using alternative methods, even when both are run using unbiased alignments. Thirdly, we have so far only tested FixVAF with sequencing data aligned with Isaac v03.16.02.19. This pipeline version was of particular interest as it is the implementation being used to align sequencing data in the 100KGP. Finally, FixVAF soft-clips reads supporting reference alleles and therefore reduces overall effective sequencing depth, potentially increasing noise in downstream analyses.

Although bioinformatic processing only accounts for around 12% of the costs of sequencing a tumor-normal sample pair (Schwarze et al., 2019), it is essential that rigorous benchmarking of bioinformatic pipelines is conducted if expensive, large-scale sequencing projects of human cancer are to achieve their true worth.

## ACKNOWLEDGEMENTS

This work was supported by grants from Cancer Research UK [grant number C1298/A8362], Myeloma UK, Bloodwise and a David Forbes Nixon Foundation Fellowship [MK]. We are grateful to the National Cancer Research Institute Haemato-oncology group and to those investigators that recruited patients to the Myeloma XI trial.

## AUTHOR CONTRIBUTIONS

AJC, DC and RSH conceived and designed the study. AJC, DC, AF, PHH and DCW provided code and performed bioinformatic analyses. MK acquired sequencing data. AJC, DC and RSH drafted the Article, which was reviewed and approved by all other authors.

